# Moment-to-moment fluctuations in neuronal excitability bias subjective perception rather than decision-making

**DOI:** 10.1101/151324

**Authors:** Luca Iemi, Niko A Busch

## Abstract

Perceiving an external stimulus not only depends on the physical features of the stimulus, but also fundamentally on the current state of neuronal excitability, indexed by the power of ongoing alpha oscillations. Recent studies suggest that heightened excitability does not improve perceptual acuity, but biases observers to report the presence of a stimulus regardless of its physical presence. It is unknown whether this bias is due to changes in observers’ subjective perceptual experience (perceptual bias) or their perception-independent decision-making strategy (decision bias). We tested these alternative interpretations in an EEG experiment in which human participants performed two-interval forced choice (2IFC) detection and discrimination. According to signal detection theory, perceptual bias only affects 2IFC detection, but not discrimination, while interval decision bias should be task-independent. We found that detection was optimal in trials in which excitability before the stimulus-present interval exceeded that before the stimulus-absent interval, consistent with an effect of excitability on perceptual bias. By contrast, discrimination accuracy was unaffected by excitability fluctuations between intervals, ruling out an effect on interval decision bias. We conclude that the current state of neuronal excitability biases the perceptual experience itself, rather than the decision process.

## Introduction

Neuronal activity just preceding, or even in the absence of experimental events, is ubiquitous in electrophysiological recordings in the form of “ongoing”, “spontaneous”, or “pre-stimulus” oscillations: rhythmic signals reflecting fluctuations of membrane potentials (Buzsaki and Draguhn, 2004). A prominent type of such spontaneous activity is the alpha rhythm (8–12 Hz), which is thought to play a key role in regulating cortical excitation and inhibition (Jensen and Mazaheri, 2010). Specifically, numerous studies have shown that states of weak *α* oscillations reflect increased excitability in sensory brain areas, as indexed by the spike-firing rate (Haegens et al., 2011), multiunit activity (van Kerkoerle et al., 2014; Becker et al., 2015), ongoing *γ* power (Spaak et al., 2012) and the hemodynamic fMRI signal (Goldman et al., 2002; Becker et al., 2011; Scheeringa et al., 2011; Harvey et al., 2013; Mayhew et al., 2013).

How do spontaneous neural oscillations interact with the processing of sensory events? A growing body of evidence demonstrates that observers are more likely to detect visual targets that are preceded by weak pre-stimulus *α*-oscillations, reflecting stronger neuronal excitability (Ergenoglu et al., 2004; van Dijk et al., 2008; Busch et al., 2009; Chaumon and Busch, 2014). But do states of strong excitability help observers see *better*, or do they simply make them see or report *more*? Recent studies have demonstrated, using signal detection theory (SDT; Green and Swets, 1966) that in visual yes/no detection, but not two-choice discrimination tasks, strong excitability increases both the hit rate and false alarm rate (Limbach and Corballis, 2016; Iemi et al., 2017; Samaha et al., 2017b). Thus, contrary to the previously dominant view in the literature (Ergenoglu et al., 2004; van Dijk et al., 2008; Romei et al., 2008; Roberts et al., 2014; Payne and Sekuler, 2014), heightened excitability evidently reflects a state of more liberal detection bias, rather than of improved detection sensitivity. This finding enables reconciling a large body of literature (see Iemi et al., 2017, for a systemic literature analysis).

These findings could be regarded as evidence refuting an effect of excitability on perception proper, showing instead an effect on subjects’ deliberate *decision bias*: a preference to report “yes, I saw the stimulus”. However, not every change in criterion implies a change in deliberate decision strategy (Witt et al., 2015); excitability might alternatively modulate *perceptual bias*: a change in the amplification of the neural representation of both signal and noise. Thus, false alarms induced by perceptual bias during states of strong excitability may be due to genuine, albeit false, impressions of seeing a stimulus (i.e. “hallucination”). The question of whether neuronal excitability reflects a decision bias or a perceptual bias remains unanswered. Importantly, decision bias and perceptual bias each lead to very different theoretical interpretations of how spontaneous brain activity is related to perception and behavior.

Unfortunately, distinguishing between decision bias and perceptual bias is impossible in classical single-interval yes/no detection tasks, but this ambiguity can be resolved in a two-interval forced choice (2IFC) detection paradigm, in which observers report, during which of two intervals the stimulus was presented. In fact, a change in “yes”-decision bias cannot influence performance in a 2IFC detection paradigm, since “yes, I saw it” is not a response option. Rather, in 2IFC paradigms, a change in decision bias would manifest as an interval bias, whereby observers would show a tendency to report the interval with stronger excitability in any perceptual task, i.e. both detection and discrimination. By contrast, changes in perceptual bias only affect 2IFC detection, but not 2IFC discrimination. In 2IFC *detection*, a perceptual bias, resulting from an imbalance of excitability between the stimulus-present and the stimulus-absent interval is expected to affect detection performance. In 2IFC *discrimination* however, a perceptual bias is expected to leave performance unaffected. This is because a shift in global excitability in one interval equally affects the response of all feature detectors, without changing their relative strength, which is the determinant of discrimination performance (see *Methods* for details).

We found that the accuracy of 2IFC detection was predicted by a state of reduced *α*- and *β*-power before the stimulus-present interval, compared to before the stimulus-absent interval. This is equivalent to a perceptual bias, resulting from a state in which excitability during the stimulus-present interval exceeds the excitability during the stimulus-absent interval. Importantly, no such effect was found for 2IFC discrimination. This rules out an effect of excitability on interval decision bias. In sum, this pattern of effects is consistent with the notion that fluctuations in excitability, indexed by *α*- and *β*-power, reflect a perceptual bias. We propose that heightened excitability, during states of weak *α*- and *β*-power, biases subjective perceptual experience, rather than strategic decision-making.

## Methods

### Signal Detection Models

In a 2IFC detection task, a faint stimulus appears in either one of two successive intervals and the task is to report which interval contained the stimulus. Signal detection theory (SDT) posits that observers sample an internal response in each interval, compare the two internal responses, and report whichever interval yielded the stronger response. Thus, if the internal response during the stimulus-present interval exceeds the response during the stimulus-absent interval, the participant makes a correct detection (difference model Green and Swets, 1966; Egan, 1975; Macmillan and Creelman, 2005; Wickens, 2002, Figure 1*a*). In a 2IFC discrimination task, each of two intervals contains a stimulus, one of which has a target feature (e.g. a specific orientation) and the task is to report which interval contained the target. SDT posits that in each interval, observers sample the internal responses of two feature detectors selective for the target and non-target stimulus, respectively. The relative strength of these two responses serves as an index of evidence of target-presence in a given interval. Observers compare this evidence between both intervals and report whichever interval yielded the strongest evidence (Green and Swets, 1966, Figure 1*b*). Note that 2IFC detection is based on comparing the responses of a single signal detector across two intervals, while 2IFC discrimination is based on comparing the relative strength of two feature detectors across two intervals.

**Figure 1:**
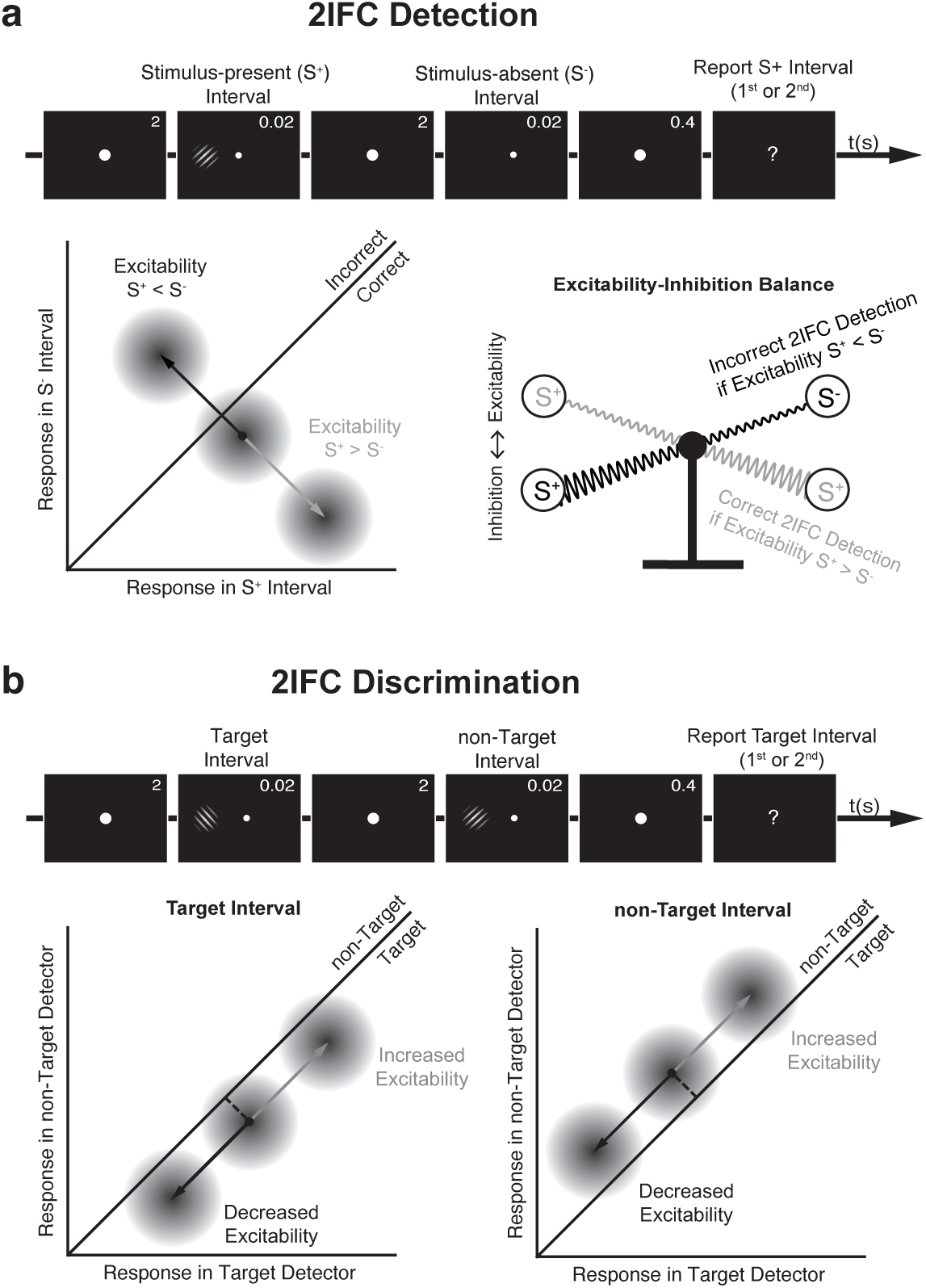
Effects of perceptual bias on 2IFC detection and discrimination performance according to signal detection theory (SDT). **a**: in 2IFC detection, a signal appears in either one of two successive intervals and the task is to report which interval contained the signal (top panel). According to SDT, the internal response in the stimulus-present interval, *RS*+, is compared to the internal response in the stimulus-absent interval, *RS*−, and the interval yielding the stronger response is reported. If *RS*+ *> RS*− (below diagonal), the report is correct, otherwise incorrect. The perceptual bias model predicts that the accuracy of 2IFC detection reports is affected by the balance of excitability and inhibition between the stimulus-present and stimulus-absent intervals (bottom panel): stronger excitability (weaker *α* and *β* power) before the stimulus-present interval, relative to before the stimulus-absent interval, is expected to boost detection accuracy; instead, stronger inhibition (stronger *α* and *β* power) before the stimulus-present interval relative to before the stimulus-absent interval is expected to impair accuracy. **b**: in 2IFC discrimination, two successive intervals contain either a target or a non-target stimulus and the task is to report in which interval the target was presented (top panel). For each interval, the difference between the internal responses of the target detector and non-target detector (i.e. distance from the diagonal; dashed line) represents the evidence of target presence in that interval, and the interval yielding stronger target-evidence is reported. If there is more target-evidence in the target interval, the response is correct, otherwise incorrect. The perceptual bias model predicts that the accuracy of 2IFC discrimination reports is not affected by increased excitability or inhibition in the target or non-target interval since both target and non-target detectors would be affected equally, leaving target evidence in that interval unchanged (bottom panel).

As explained in the Introduction, this study was designed to compare and contrast two alternative models, in which fluctuations of excitability reflect either a perceptual bias or a decision bias. In short, a perceptual bias would make anything (signal as well as noise) look more like a stimulus, while a decision bias would make the observer more inclined to report that there was a stimulus. Additionally, we tested a sensitivity model, in which increased excitability improves the accuracy of discrimination (Payne and Sekuler, 2014), such as discerning stimulus from no-stimulus and target from non-target. This sensitivity model was tested to replicate, using a 2IFC paradigm, three recent studies reporting a null effect of excitability on single-interval yes/no detection sensitivity (Limbach and Corballis, 2016; Iemi et al., 2017) and 2-alternative-forced-choice (2AFC) discrimination accuracy (Iemi et al., 2017; Samaha et al., 2017b).

According to the perceptual bias model, heightened neuronal excitability, indexed by reduced pre-stimulus oscillations, amplifies internal responses to both signal and noise. Hence, 2IFC detection accuracy should be influenced by the balance of excitability and inhibition between the stimulus-present and stimulus-absent interval. Specifically, stronger excitability (reduced pre-stimulus oscillations) in the stimulus-present interval would amplify the internal response to the signal, while stronger inhibition in the stimulus-absent interval (increased pre-stimulus oscillations) would reduce the internal response to the noise. The net effect of this beneficial balance would be to boost the separability of the signal and noise representations, resulting in more accurate detections. Conversely, relatively stronger inhibition in the stimulus-present interval, and relatively stronger excitability in the stimulus-absent interval, would make the signal less discernible and thus impair detection accuracy (see Figure 1*a*). By contrast, the perceptual bias model predicts no effect on 2IFC discrimination accuracy. Since any change in excitability in one interval affects both target and non-target feature detectors equally, the relative strength of their responses (i.e. target-evidence in this interval) remains unchanged (see Figure 1*b*).

According to the decision bias model, fluctuations in excitability influence observers’ strategic decision behavior rather than their perceptual processes. Note that a bias to report “yes, I saw it” – as in a single interval yes/no detection task – cannot influence decisions in a 2IFC detection task. Rather, a decision bias would manifest itself as an interval bias, whereby observers would display a tendency to report the interval with stronger excitability. Crucially, this interval decision bias should apply to both 2IFC detection and 2IFC discrimination.

According to the sensitivity model, fluctuations in excitability reflect changes in perceptual acuity. Thus, the sensitivity model predicts that accuracy is related to the overall power in both intervals, not to the difference between intervals as predicted by the bias models. Specifically, correct 2IFC detection and discrimination should be associated with increased excitability, reflected by reduced pre-stimulus oscillations in both intervals.

### Participants

Previous studies on the relationship between neuronal excitability and perception (Busch et al., 2009; Lange et al., 2013; Chaumon and Busch, 2014; Iemi et al., 2017, e.g.) have typically reported samples of 12-33 participants. To ensure a robust estimate of our neurophysiological effect, we recruited a sample of 25 participants (mean: 29.3, SEM = 0.75 year old, 16 females, 2 left-handed). All participants had normal or corrected-to-normal vision and no history of neurological disorders. Each participant took part in two sessions, one for each task, on two separate days within a 7-day period. One participant was excluded before EEG preprocessing and behavioral analysis, because she could not participate in the second experiment. Two participants were excluded after EEG preprocessing because of excessive artifacts. A total of twenty-two participants were included in the analysis. Prior to the experiment, written informed consent was obtained from all participants. All experimental procedures were approved by the ethics committee of the German Psychological Society.

### Stimuli

Both experiments were written in MATLAB (RRID:SCR_001622) using the Psychophysics toolbox 3 (RRID:SCR_002881; Brainard, 1997; Pelli, 1997). Stimuli were presented on a black background, using a gamma-linearized cathode ray tube monitor operated at 100 Hz and situated in a dark room. Low-contrast Gabor patches tilted by 10° clockwise or counterclockwise from the vertical meridian with a diameter of 0.75° visual angle were displayed at 10° to the left or to the right of the fixation cross.

Each trial included two successive intervals, separated by a 2 s gap. Each interval lasted two frames (0.02 s) and was indicated by a 50% reduction in the diameter of the fixation point (Figure 1). A Gabor stimulus was presented in one or both intervals for a duration of two frames (0.02 s). After a delay of 400 ms following the second interval, the fixation dot turned into a question mark, which instructed the participants to deliver a response via button-pressing, in accordance with the task instructions. At the end of each trial, participants received color-coded feedback. The fixation dot was then displayed again and a new trial started.

### Experimental Design

Participants performed a 2IFC detection and a 2IFC discrimination task in two separate sessions. In the 2IFC detection task, the target stimulus was presented in either one of two successive temporal intervals and a blank screen in the other interval (Figure 1). Participants were informed that each trial contained a stimulus, which could appear in either interval, and were instructed to report in which interval they perceived the stimulus (“first” vs. “second”). In the 2IFC discrimination task, a left-tilted and right-tilted stimulus were presented in two successive temporal intervals. Participants were informed that each trial contained two stimuli characterized by different tilts, and were instructed to report in which interval they perceived the left-tilted target stimulus (“first” vs. “second”). For both tasks, we denote trials as S1 or S2 trials, according to the interval in which the stimulus occurred in the 2IFC detection, or the target stimulus occurred in the 2IFC discrimination. Stimulus side, tilt, and target interval were counterbalanced across trials in both tasks. Task order was counterbalanced across participants. For each participant and task, an adaptive staircase procedure (QUEST Watson and Pelli, 1983) was used to find a stimulus contrast yielding 75% accuracy. To ensure that the analysis included only trials of similar contrast, we rejected outlier trials in which the difference between the presented contrast value and the final threshold estimated by QUEST, exceeded a criterion value. One analysis used a conservative criterion of ±0.04 contrast units for trial rejection, yielding a homogeneous range of contrast values. However, using this conservative criterion (see next paragraph), we noticed that more trials were rejected for the discrimination task than for the detection task. Thus, we repeated the analysis using a more liberal criterion of ±0.2 contrast units, which yielded a comparable number of trials in both tasks. Each session lasted approximately 1,5 h with breaks and included 700 trials divided into 14 blocks of 50 trials each.

### EEG recording and preprocessing

EEG was recorded with a 64-channel Biosemi ActiveTwo system at a sampling rate of 1024 Hz. Electrodes were placed according to the international 10-10 system. The horizontal and vertical electro-oculograms were recorded by additional electrodes at the lateral canthi of both eyes and below the eyes, respectively.

The EEGLAB toolbox version 11, running on MATLAB (R2010b), was used to process and analyze the data (Delorme, 2004). Data were re-referenced to the averaged mastoids, epoched from -3700 to 700 ms relative to the second interval onset and down-sampled to 256 Hz. The data were then filtered using an acausal band-pass filter between 0.25 and 80 Hz. Gross artifacts (eye blinks, and noisy data segments) were removed manually, and entire trials were discarded when a blink occurred within a critical 0.5 s time window preceding interval onset, to ensure that participants’ eyes were open at interval onset. After rejecting trials with EEG artifacts and contrast outliers, the total number of trials analyzed for the detection session was 675 (5.50 SEM) and 681 (5.49 SEM), using the conservative and liberal contrast criterion, respectively. For the discrimination session, the total number of trials analyzed was 546 (27.36 SEM) and 662 (8.22 SEM), using the conservative and liberal contrast criterion, respectively.

Noisy channels were selected manually for interpolation with the data from the adjacent channels. Furthermore, the EEG data were transformed using independent component analysis (ICA), and SASICA (Chaumon et al., 2015) was used to guide the exclusion of IC related to noisy channels and muscular contractions, as well as blinks and eye movements occurring before or after the intervals. For the time-frequency analysis, we re-epoched the trials relative to the onset of a single interval, allowing us to analyze between-interval fluctuations in excitability. Time-frequency analysis was carried out using a wavelet transform (Morlet wavelets, 30 frequencies, frequency range: 5-30 Hz, number of cycles increasing linearly from 1 to 12). Thus, a wavelet at 10 Hz was 4.4 cycles long and had a temporal resolution *σ_t_* of 0.14 s and a spectral resolution *σ_f_* of 4.53 Hz. Since wavelet analysis is computed by convolving the data with a function that is extended in time, it is possible that pre-stimulus effects close to stimulus onset are actually affected by poststimulus data. Iemi et al. (2017) determined the extent of this contamination by estimating the wavelet’s temporal resolution *σ_t_* (Tallon-Baudry et al., 1996). Thus, we consider effects as truly “pre-stimulus”, only if they occur at time points earlier than interval onset - *σ_t_*. We indicate this time limit with a red line in Figure 2*a/c*.

**Figure 2:**
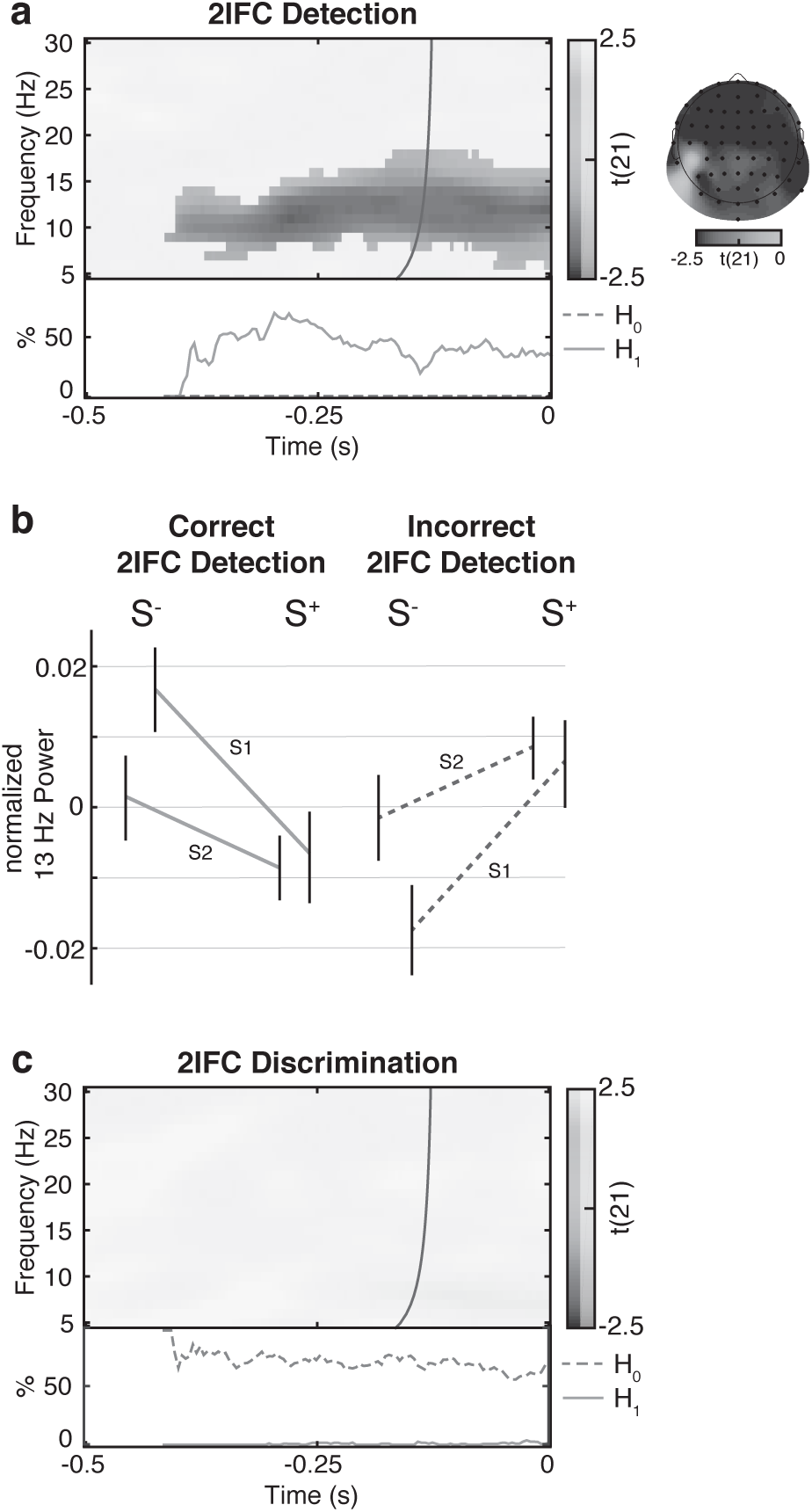
Effect of between-interval fluctuations of oscillatory power on perceptual accuracy. **a**: Group-level t-statistics map of the regression coefficient *β*_1_, indicating the effect of power differences between the window before the stimulus-present interval and that before the stimulus-absent interval on 2IFC detection accuracy. Reduced *α* and *β* power preceding the stimulus-present interval relative to that preceding the stimulus-absent interval predicts correct performance. **b**: Group-average pre-interval power for correct and incorrect trials, normalized by the power in all trials. Results are shown for illustrative purposes for the time window, frequency, and electrode that yielded the strongest cluster-level effect (-.250 ms, 13 Hz, FCz). Correct detection is associated with stronger pre-stimulus power in the stimulus-absent interval (S-) and weaker power in the stimulus-present interval (S+), while incorrect detection is associated with the opposite pattern. This effect appears regardless of whether the stimulus is in interval 1 (S1) or in interval 2 (S2). **c**: Group-level t-statistics map of the regression coefficient *β*1, indicating the effect of power differences between the window before the target interval and that before the non-target interval on 2IFC discrimination accuracy. Between-interval fluctuations of power power do not predict 2IFC discrimination accuracy. This null effect is corroborated by the BF analysis, indicating that there is more evidence supporting the *H*0 than supporting the *H*1. The maps in **a** and **c** are averaged across the electrodes comprising the detection cluster and masked by a p-value of 0.05 using two-sided cluster permutation testing. Time 0 ms indicates interval onset. The bold line in **a** and **c** indicates the time points before which oscillatory activity is not contaminated by stimulus-related activity (Iemi et al., 2017).

### Statistical analysis

#### Generalized linear modeling

The predictions of the bias models concern the *relative* magnitude of excitability between the two intervals within a trial. Thus, for each single trial, time point, frequency, and electrode, we computed a measure, *P_rel_*, comparing oscillatory power between the two intervals as:

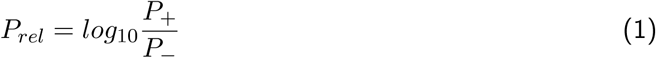

where *P*_+_ is power in the interval containing the stimulus in 2IFC detection, and the stimulus with the target feature in 2IFC discrimination and *P*_−_ is power in the stimulus-absent interval in 2IFC detection and the stimulus with the non-target feature in 2IFC discrimination. For example, values of *P_rel_ >* 0 at time *t* = 0 in the 2IFC detection task indicate that power was stronger at the onset of the stimulus-present interval, compared to power at the onset of the stimulus-absent interval.

The predictions of the sensitivity model concern, instead, the *average* magnitude of excitability in the two intervals within a trial. Thus, for each single trial, time point, frequency, and electrode, we computed a measure, *P_ave_*, reflecting the average power in the two intervals within a trial.

*P_rel_* and *P_ave_* were analyzed for frequencies between 5 and 30 Hz and between -500 and 0 ms relative to interval onset. S1 and S2 trials were collapsed for this analysis.

Next, we modeled the effects of *P_rel_* and *P_ave_* on correct/incorrect reports using generalized linear modeling (GLM). For each participant and for each electrode, frequency, and time point, we fit a regression model of the form:

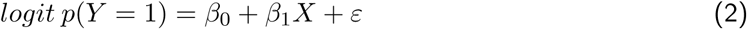

where *p*(*Y* = 1) is the probability of a correct response, *Y* the correct/incorrect response variable dummy-coded as a 1/0 variable, X the predictor variable (*P_rel_* or *P_ave_*), *β*_0_ and *β*_1_ the estimated coefficients, and *ε* the residual errors of the GLM. This corresponds to a logistic regression model, where the coefficient *β*_1_ represents the contribution of the predictor (*P_rel_* or *P_ave_*) on the probability of correct response. GLMs were fit separately for detection and discrimination tasks.

#### Group-level statistical testing

We then tested whether the regression coefficients *β*_1_ at each electrode, frequency, and time point were significantly different from 0 across participants, by computing the t-statistics of these regression coefficients against the null hypothesis that there are no effects (*β*_1_ = 0). To correct for multiple comparisons, we used a nonparametric cluster permutation test (Maris and Oostenveld, 2007) on the absolute values of the t statistics with 1000 permutations, a cluster threshold p-value of 0.05, and a final significance p-value of 0.05. This test was two-sided. Again, this procedure was computed separately for the detection and discrimination tasks.

#### Bayes factor analysis

To provide evidence supporting the perceptual bias model (Figure 1), we need to demonstrate an effect of between-interval fluctuations of excitability on 2IFC detection performance, but a null-effect on 2IFC discrimination performance. However, in conventional inferential statistics, a non-significant result only indicates that the null hypothesis cannot be rejected; it does not necessarily follow that the null hypothesis is actually true. It is possible that the data might be inconclusive, e.g. due to insufficient statistical power. Thus, to directly estimate evidence supporting the null hypothesis, we used Bayes factor analysis (Rouder et al., 2009). We first estimated the JZS Bayes factor (BF) for the negative t-statistics of the difference maps, setting the prior on effect size following a Cauchy distribution with a scale factor 0.707, as recommended by Rouder et al. (2009). Conventionally, BF indicates whether there is evidence supporting the alternative hypothesis (*H*_1_: *β*_1_ = 0, if *BF >* 3) or supporting the null hypothesis (*H*_0_: *β*_1_ = 0, if *BF <* 1*/*3) or whether the evidence is inconclusive (if 1*/*3 *< BF <* 3). We then computed for each time point, the proportion of electrodes and frequencies yielding evidence for *H*1 and *H*0 (see insets below the time-frequency plots in figure 2*a/c*).

Note that throughout the text, the term “effect” is used to indicate a significant statistical outcome, but not a causal relationship. To determine whether the nature of the observed effect is correlative or causal, other approaches, such as neurostimulation (Helfrich et al., 2014; Romei et al., 2010), are necessary.

## Results

### Behavior

For each participant, an adaptive staircase procedure (see *Methods*) adjusted stimulus contrast to ensure a proportion of correct responses of 75% in both the 2IFC detection and the discrimination task. The participants included in the analysis had a mean proportion of correct detection responses of 73.2 % (SEM = .002) and a mean proportion of correct discrimination responses of 72.5 % (SEM = .007), indicating that the staircase procedure was successful. On average, the stimulus contrast necessary for achieving this level of performance was higher in the 2IFC discrimination than in the 2IFC detection task (two-tailed paired-sample t test: t(21) = 5.77, p < 0.001), which is consistent with previous work (Iemi et al., 2017).

### EEG

#### Perceptual bias vs. Decision Bias Models

To test the predictions of perceptual and decision bias models, we modeled the effects of between-interval fluctuations of oscillatory power (as a measure of excitability) on 2IFC detection and discrimination accuracy, using a GLM approach. For 2IFC detection, the analysis using a conservative trial rejection criterion (see *Methods*) yielded a significant negative cluster starting from -.414 ms before interval onset, at frequencies between 6 and 18 Hz (Figure 2a). In other words, weak *α* and *β* power in the time window before the stimulus-present interval (relative to the stimulus-absent interval) predicted correct detection. The topography of this effect was widespread. The peak of this cluster was at electrode FCz, at 13 Hz and at -.250 ms before interval onset (t(21) = -5.90) (see Figure 2*a*).

No significant cluster was found for 2IFC discrimination using either a conservative (Figure 2b) or liberal trial rejection criterion (results not shown). In other words, differences in oscillatory power between the window preceding the target interval and that preceding the non-target interval did not predict discrimination accuracy.

To validate these findings and to provide evidence of a true null effect on 2IFC discrimination accuracy, as opposed to merely inconclusive evidence, we quantified, for each time point, the proportion of cluster frequencies and electrodes at which the data provided evidence of an effect (*H*_1_) or evidence of a null effect (*H*_0_). For the time points, frequencies and electrodes within the cluster of significant effects in 2IFC detection, the proportion of data points in favor of an effect on perceptual accuracy, by far outnumbered the proportion of data points in favor of a null effect (*H*_1_ *> H*_0_, bottom inset of Figure 2*a*). This result is expected, because the BF analysis was restricted to the cluster with significant effects as determined by the cluster permutation test. However, our primary interest was to compare the strength of evidence in favor of an effect on 2IFC detection accuracy to the strength of evidence against an effect on 2IFC discrimination accuracy. Since no significant cluster of effects was found for 2IFC discrimination, we conducted this comparison for the time points, frequencies, and electrodes within the detection cluster, following a method used in Iemi et al. (2017). Note that the decision bias model predicts the same effect of between-interval power fluctuations for 2IFC detection and 2IFC discrimination, so that effects should coincide in time, frequency, and space for the two tasks. The proportion of data points in favor of a null effect on discrimination accuracy by far outnumbered the proportion of data points in favor of an effect (*H*_0_ *> H*_1_, bottom inset of Figure 2c). This analysis was performed using the conservative rejection criterion. Since more contrast outlier trials were rejected for the discrimination task using the conservative rejection criterion, we repeated this analysis using a more liberal contrast rejection criterion, which yielded a similar number of trials in both tasks (see *Methods*). This analysis yielded qualitatively similar results (results not shown), indicating that this finding was not due to a lower number of discrimination trials. In sum, the effect of between-interval fluctuations of oscillatory power on 2IFC discrimination accuracy was not merely weak or inconclusive, but entirely absent.

#### Sensitivity Model

To test the predictions of the sensitivity model, we modeled the effects of the averaged power across the intervals (as a measure of excitability) on 2IFC detection and discrimination accuracy, using a GLM approach. No significant cluster was found, neither for 2IFC detection nor discrimination, using either a conservative or liberal trial rejection criterion (results not shown). We then tested whether there was evidence of a null effect of average oscillatory power on perceptual accuracy, using Bayes factor analysis, as described above. The proportion of data points in favor of a null effect on both 2IFC detection and discrimination accuracy by far outnumbered the proportion of data points in favor of an effect.

## Discussion

### Excitability modulates perceptual bias rather than decision bias

What are the perceptual consequences of spontaneous fluctuations in neuronal excitability? Accumulating evidence suggests that, during states of strong neuronal excitability, indexed by weak *α* and *β* ongoing oscillations, observers are more likely to report the presence of a sensory stimulus, irrespective of its actual physical presence. Thus, contrary to the previously dominant view (e.g. Ergenoglu et al., 2004; Payne and Sekuler, 2014), strong excitability reflects a state of liberal detection criterion/bias rather than of improved perceptual acuity/sensitivity. What is the mechanism linking fluctuations of excitability and bias? According to SDT, two alternative mechanisms are possible. On the one hand, strong excitability could indicate a state of more permissive detection strategy, during which observers prefer to report “yes I saw the stimulus”. This mechanism is referred to as *decision bias*. On the other hand, strong excitability could reflect a state of increased baseline sensory processing, resulting in an amplification of the neural responses to both signal and noise. At the behavioral level, this is paralleled by an amplification of subjective perceptual experience, during which observers “perceive” stimuli even when they are not physically present. This mechanism is referred to as *perceptual bias*. Past studies using single-interval detection tasks (Iemi et al., 2017; Limbach and Corballis, 2016) were unable to distinguish between these alternative interpretations of excitability, because SDT cannot determine the underlying source of the bias, be it perceptual or decision-based (Wixted and Stretch, 2000; Witt et al., 2015).

In this study, we addressed the issue by analyzing the effects of pre-stimulus oscillations, as a measure of excitability, on 2IFC detection and discrimination tasks, which are differently affected by perceptual bias and decision bias. The predictions of an SDT model of perceptual bias are twofold. First, 2IFC detections should be more accurate, when excitability in the signal interval exceeds that in the no-signal interval, due to an amplification of the signal representation and dampening of the noise representation. Second, in 2IFC discrimination, fluctuations of excitability between target and non-target intervals should not affect discrimination accuracy. This is because a change in global excitability (i.e. not specific to a certain feature value) affects the response of all feature detectors equally, without changing their relative strength, on which discrimination accuracy depends. By contrast, an SDT model of decision bias predicts that fluctuations in excitability influence the observer’s strategic decision behavior, rather than perceptual processing. Note that a “yes”-bias, as in a single-interval detection task, cannot affect decisions in a 2IFC detection task, because “yes, I saw it” is not among the given options. However, an interval decision bias predicts a tendency to report the interval with stronger excitability, regardless of perceptual task, and should therefore be manifest in both 2IFC detection and 2IFC discrimination.

To test these alternative models, we predicted 2IFC detection accuracy, based on a comparison of excitability between stimulus-present and stimulus-absent intervals, and 2IFC discrimination accuracy based on a comparison of excitability between target- and non-target intervals. We found that detection accuracy was highest when pre-stimulus *α*- and *β*-power was lower before the stimulus-present interval relative to the stimulus-absent interval. This effect rules out a “yes”-decision-bias that might have affected previous findings from single-interval detection tasks (e.g. Chaumon and Busch, 2014; Limbach and Corballis, 2016; Iemi et al., 2017). Moreover, we found evidence that discrimination accuracy was unaffected by between-interval fluctuations of excitability, ruling out an interval decision bias model. Taken together, the effect on 2IFC detection and the evidence of a null effect on 2IFC discrimination confirm the predictions of the perceptual bias model.

### Excitability does not affect perceptual acuity

It is important to note that the effect of perceptual bias on 2IFC detection, i.e. when excitability is specifically strong in the stimulus-present interval, merely represents a serendipitous distortion of subjective perception “in the right direction” rather than an actual improvement in perceptual acuity. By contrast, a sensitivity model predicts that excitability in both intervals improves detection and discrimination. However, we found that excitability, averaged across both intervals, affected neither 2IFC detection nor discrimination accuracy. This result replicates, in a 2IFC paradigm, the findings of three recent studies, reporting a null effect of excitability on single interval yes/no detection sensitivity (Limbach and Corballis, 2016; Iemi et al., 2017) and 2AFC discrimination accuracy (Iemi et al., 2017; Samaha et al., 2017b). Taken together, these findings challenge the long-held notion that neuronal excitability affects the accuracy of perceptual decisions. This notion has been based on the observation that successful stimulus detection (hit-rate) is associated with relatively stronger excitability. Such findings have been obtained with visual (Ergenoglu et al., 2004; van Dijk et al., 2008), auditory (Leske et al., 2015) and somatosensory detection (Baumgarten et al., 2016) and for detection of TMS-induced phosphenes (Romei et al., 2008; Samaha et al., 2017a). However, without testing for an effect on the false-alarm rate on stimulus-absent trials, it is possible that excitability affects rather the bias to report a stimulus, irrespective of whether or not this is accurate. Indeed, recent studies analyzing signal detection measures found that increased excitability is associated with a more liberal detection bias in both vision (Limbach and Corballis, 2016; Iemi et al., 2017) and somatosensation (Craddock et al., 2017). Our results are also consistent with several experiments that found no effect of excitability on multiple-alternative-forced-choice (mAFC) *discrimination* performance (e.g. Bays et al., 2015; see Iemi et al., 2017 for a comprehensive literature review). An SDT model of perceptual bias, in fact, predicts these null findings because discrimination performance in mAFC tasks is unaffected by a “yes”-bias, and a modulation of excitability does not change the discriminability between response alternatives (Iemi et al., 2017).

Samaha et al. (2017b) replicated the finding that states of high excitability do not improve 2AFC accuracy, and additionally demonstrated that excitability instead biases observers to report higher confidence in their 2AFC decisions. An SDT model of perceptual bias predicts this finding because confidence, unlike accuracy, is thought to depend on the absolute amount of evidence in favor of a perceptual choice, be it accurate or not, regardless of the amount of evidence against this choice (Zylberberg et al., 2012; Maniscalco et al., 2016). According to a perceptual bias model, a state of increased excitability is expected to boost the representation of both stimulus alternatives, yielding increased confidence reports regardless of the accuracy of the perceptual decision.

### Within-trial fluctuations of excitability

Past studies on the relationship between excitability and perception have typically analyzed the relationship between behavioral and neuronal measures on a trial-by-trial basis: i.e. differences in the hit rate throughout the experiment were related to differences in *α* power *across* trials (e.g. Busch et al., 2009; Iemi et al., 2017; Limbach and Corballis, 2016). This across-trial approach treats individual trials as independent samples and therefore ignores the fact that data are collected in temporal order. This is potentially problematic, because it is known that both detection rate and excitability change over the course of an experiment. Specifically, behavioral measures such as hit rate (Boncompte et al., 2016; Carrasco-López et al., 2017) and sensitivity (Maniscalco et al., 2017) tend to decrease over time, possibly due to progressive fatigue, resulting from an exhaustion of cognitive resources. Likewise, ongoing *α*-power tends to increase over the course of an experiment (van Dijk et al., 2008), suggesting a decrease in excitability, possibly as a result of fatigue (Kaida et al., 2006). Therefore it is possible that the across-trial correlation between behavioral and neuronal measures in previous studies may be a by-product/epiphenomenon related to fatigue.

To test the bias models in our study, we used a different approach and quantified on a trial-by-trial basis, the changes in excitability between two intervals within a trial lasting a few seconds, instead of the absolute magnitude of excitability within a trial. This approach ensures that our measure of excitability is not influenced by fatigue-related effects occurring over longer time scales. Crucially, our results show a significant across-trial correlation between excitability fluctuations and perceptual reports, even when the effects of fatigue are ruled out. This study thus demonstrates that the relationship between excitability and perception is not determined by fatigue.

### Conclusions

We propose that the current state of neuronal excitability—indexed by spontaneous neural oscillations—biases the observer’s subjective perceptual experience, by amplifying or attenuating sensory signals and noise, rather than the decision strategy.

## Author contributions statement

N.A.B and L.I conceived the experiment, L.I. conducted the experiment, L.I. analyzed the results. Both authors wrote and reviewed the manuscript.

## Additional information

No financial interests or conflicts of interest declared

## Acknowledgments

We thank Alberto Mariola and Joseph Wooldridge for assistance with the EEG recording and preprocessing, as well as Jason Samaha and Esra Al for useful discussions. This work was supported by the German Academic Exchange Service DAAD (LI), the Berlin School of Mind and Brain (LI), and National Research University Higher School of Economics (LI). The authors are grateful to Dr. Brian Bloch for his editing of the manuscript.

